# Turning vice into virtue: Using Batch-Effects to Detect Errors in Large Genomic Datasets

**DOI:** 10.1101/189670

**Authors:** Fabrizio Mafessoni, Rashmi B Prasad, Leif Groop, Ola Hansson, Kay Prüfer

## Abstract

It is often unavoidable to combine data from different sequencing centers or sequencing platforms when compiling datasets with a large number of individuals. However, the different data are likely to contain specific systematic errors that will appear as SNPs. Here, we devise a method to detect systematic errors in combined datasets. To measure quality differences between individual genomes, we study pairs of variants that reside on different chromosomes and co-occur in individuals. The abundance of these pairs of variants in different genomes is then used to detect systematic errors due to batch effects. Applying our method to the 1000 Genomes dataset, we find that coding regions are enriched for errors, where about 1% of the higher-frequency variants are predicted to be erroneous, whereas errors outside of coding regions are much rarer (<0.001%). As expected, predicted errors are less often found than other variants in a dataset that was generated with a different sequencing technology, indicating that many of the candidates are indeed errors. However, predicted 1000 Genomes errors are also found in other large datasets; our observation is thus not specific to the 1000 Genomes dataset. Our results show that batch effects can be turned into a virtue by using the resulting variation in large scale datasets to detect systematic errors.

## Introduction

Next generation sequencing technologies allowed for the generation of datasets that include genetic data of a large number of individuals. To produce these datasets, sequencing data of different coverage and from different platforms or different batches of sequencing chemistry may need to be combined. This can result in differences in the types of errors and different amounts of errors across samples (Schirmer, et al. 2015; Torkamaneh, et al. 2016; Wall, et al. 2014; Wolpin, et al. 2014).

Here, we introduce a method to identify individual genomes with a higher error rate in large datasets and to predict which variants are likely due to error. The method first tests pairs of variants that reside on different chromosomes for signals of linkage disequilibrium. Linkage between separate chromosomes is not expected by population genetics theory for a randomly mating population, unless strong epistatic interactions are present. However, such signals can occur if errors affect individual genomes differently, leading to co-occurring erroneous variants on different chromosomes in the same individuals (fig. 1). This first step of testing pairs is computationally intensive and we therefore limited the computation of linkage to pairs of variants in a subset of the genome. In the second step, we compare the contribution of individual genomes to the total linkage signal to identify outlier individuals that carry more potentially erroneous variants. As a last step, we use the differences in the number of linked pairs between individuals to test which variants are present primarily in those individuals that carry more predicted errors (fig. 1). This last step can be applied to all variants, not only those that have been tested for linkage, resulting in a list of predicted erroneous variants for the complete dataset. Removing these errors, we repeat the procedure starting from the second step until no significant differences in the burden of predicted errors is observed between individuals. No knowledge of differences in sequencing technologies or other factors are required by this approach.

**Figure 1.**
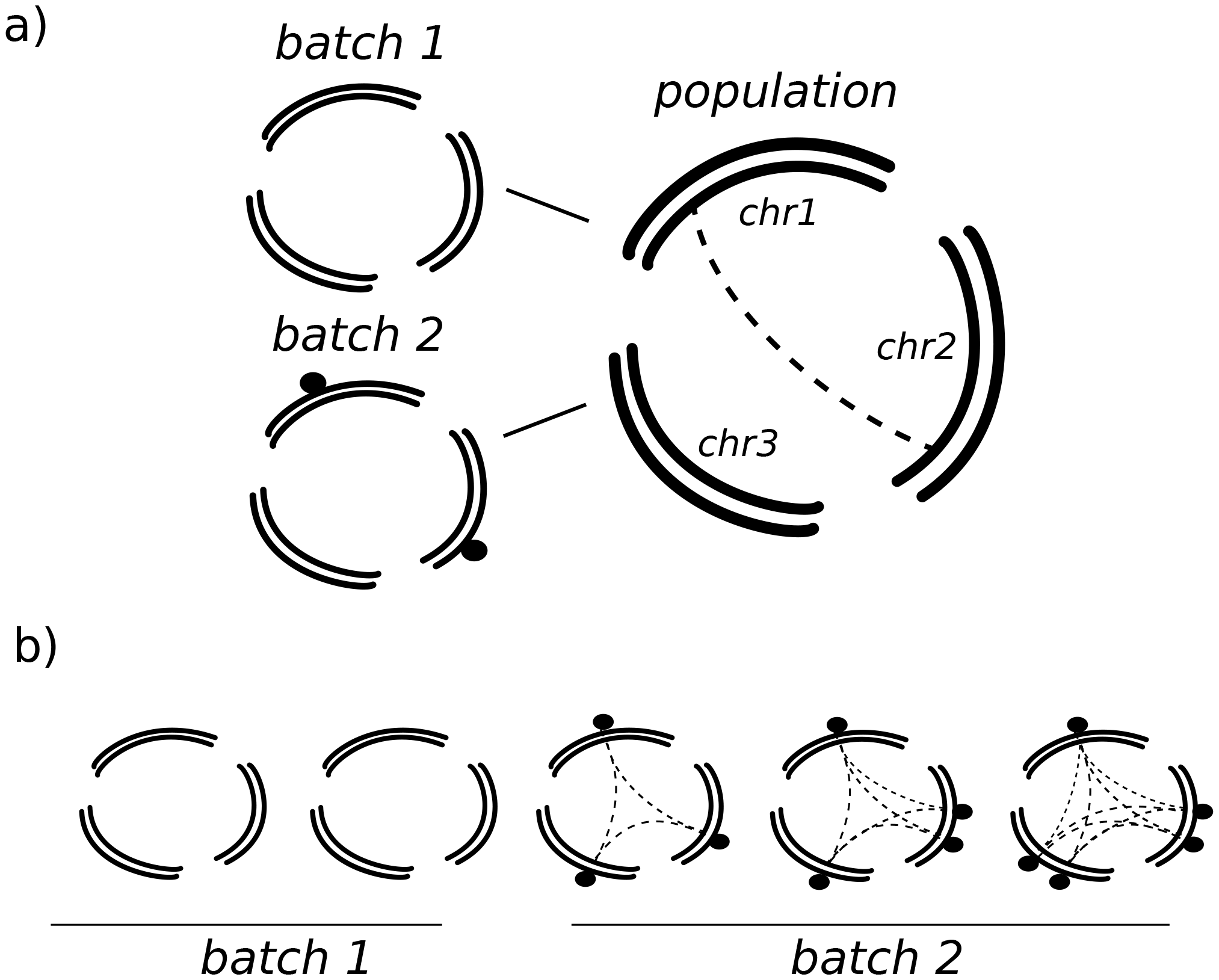
Outline of the method. a) Sequencing data generated from samples with different sequencing quality or processing might introduce different errors (black dots). Since these errors will be present in samples coming from the same platform, they will give a signal of linkage between different chromosomes (dashed lines). b) The contribution to the linkage signal can be computed for each sample (dashed lines), and used to identify samples coming from the same batch and with similar error profiles, as well as the errors. See also supplementary figure 1.

## Results

### Excess of inter-chromosomal linkage-disequilibrium in the 1000 Genomes dataset

We applied our method to the widely used 1000 Genomes dataset (Sudmant, et al. 2015; The 1000 Genomes Project Consortium 2015). The 1000 Genomes data have been acquired over 7 years, involving 10 sequencing centers, 5 sequencing technologies, and several platform versions (The 1000 Genomes Project Consortium 2015, 2012). Individuals also differ in genome-wide sequencing coverage and in the coverage of the additional exome sequencing data. We limited our analysis to populations with at least 95 unrelated individuals, resulting in a total of 12 populations that we were able to test. Since many individuals from the 12 populations contained data generated via exome capture, we considered for our analysis all rare (minor allele frequency MAF >1% and <5%) and common variants (MAF>5%) in coding regions (“coding region dataset”; 107087 sites over all 12 populations) and, as a separate dataset (“intergenic dataset”), an equal number of rare and common intergenic variants. To minimize the influence of population substructure on our measure, we calculate inter-chromosomal linkage for each population independently.

For both the intergenic and coding region datasets and for all populations, we observe an excess of linked pairs over the expected number at a false discovery rate of 5% or when comparing to an expectation derived from randomly assigning chromosomes to individuals (fig. 2a, supplementary figs. 2-3). Analyzing each population separately, we find that linked pairs are often shared between populations, but this sharing does not reflect population relationships (fig. 2b). However, much more significant links are discovered in the coding region dataset compared to the intergenic dataset. In coding regions we find that variants are often linked to other variants on several different chromosomes, leading to large clusters of paired-variants (supplementary figure 4). Maintaining such large clusters would require implausible selection pressures that favor the co-inheritance of minor alleles (supplementary figures 5). This contrasts with the concept of synergistic epistatic interaction among deleterious variants, that would lead to a repulsion between rare variants (Sohail, et al. 2017).

**Figure 2.**
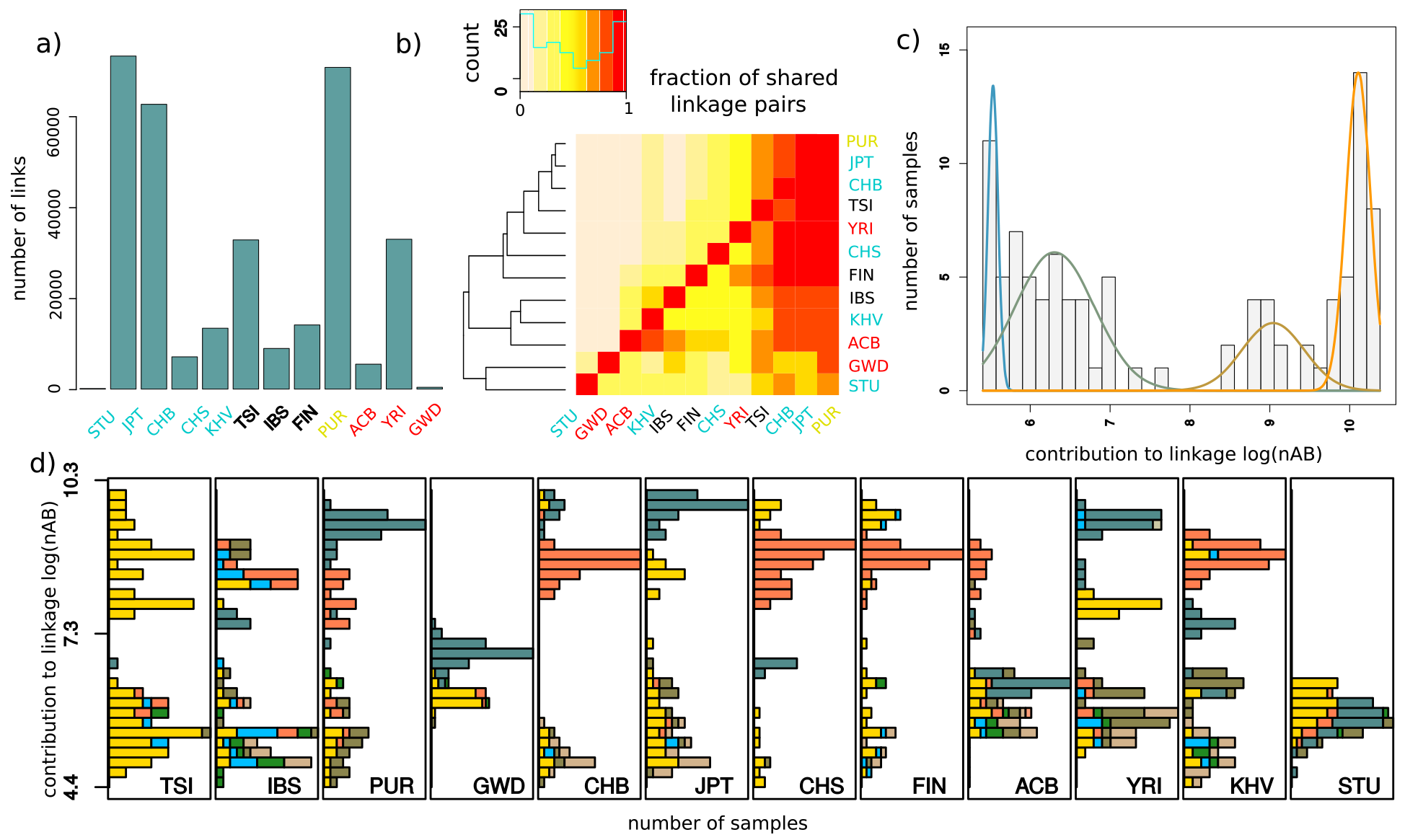
Characteristics of inter-chromosomal linkage among common variants in the coding regions (>5% minor allele frequency). a) Number of inter-chromosomal linked variants with a false discovery rate (FDR) < 5% in the 1000 Genomes populations, when analyzed independently. The FDR was calculated by comparing the p-value of each linked pair to the distribution of p-values after permuting chromosomes across individuals. Populations labels are colored according to the continent: blue for Asia, red for Africa, black for Europe and yellow for others. b) Fraction of inter-chromosomal linked variants in one population (row) that are also linked in another population (column). Darker colors indicate a higher proportion of linked variants. The order of the populations is determined by the hierarchical clustering graph shown on the left, calculated on the basis of the sharing of linked variants. c) Contribution of Chinese from Bejing individuals to the linkage signal (bars) given by the number of linked minor alleles (nAB). Individuals with similar nAB values were grouped by a Gaussian mixture model, whose fitted distributions are shown as colored lines. d) Distribution of nAB for individuals from different 1000 Genomes populations. Colors indicate the sequencing center per individual. Individuals sequenced in multiple centers were marked with a separate colors.

Next, we calculated the contribution of each individual to the overall signal of linkage in a population by summing over the estimated number of linked pairs of minor alleles this individual carries (called nAB) (Schaid 2004; Weir 1996). For this, we considered all linked pairs that showed a significant combined p-value across all populations (see Methods). We then compared the distribution of nAB over the individuals in a given population to the distribution calculated after randomly assigning chromosomes to individuals. We found that the variance in nAB is over 80-fold higher in intergenic regions and over 100-fold higher in coding regions compared to the expectation from randomization (Wilcoxon-rank test: intergenic p-value<7.4×10^−7^, coding region p-value<10^−20^), showing that the signal of linkage is driven by individuals that carry an excess of linked pairs. Interestingly, in coding regions most populations show groups of individuals that have similar nAB values, but differ from the values observed for individuals of other groups (fig 2c,d, supplementary figures 6-7). Consistent with this observation, the nAB distribution in all populations fit a model of a mixture of several Gaussian distributions significantly better than a model with just one Gaussian distribution, and we use the fitted Gaussians to assign individuals to groups (fig. 2c, supplementary figures 6-7).

### Identification of errors

To explain the differences in nAB between individuals for the coding region dataset, we correlated the group assignment of individuals from all populations with technical features of individuals annotated as part of the 1000 Genomes dataset. We found that both coverage, the presence of SNP array data for the sample and sequencing center are significantly associated with differences in nAB (coverage: p-value<10^−18^; sequencing center: p-value<10^−20^). The factor that has the strongest effects is sequencing center, explaining over 80% of the variation in nAB (fig. 2d, supplementary table 2, supplementary figure 8-9). We notice that alternative explanations are incompatible with the observed signal: for example, the possibility that polymorphic genetic rearrangements contribute to a large extent to the linkage signal is incompatible with the clustering of individuals according to their nAB, whereas population substructure would not generate the same linked variants across different populations. For the intergenic dataset, we find that sequencing center is still strongly associated with nAB, but coverage is only marginally associated when considering a minor allele frequency of 1% (supplementary table 3). In total across all populations much fewer variants appear linked in the intergenic compared to the coding region dataset (421 or ~1% of the number of linked pairs in the coding regions; see fig. 2a versus supplementary figure 10c). A larger amount of intergenic variants to determine linked pairs does not change this result..

We next searched for variants where a minor allele is preferentially encountered in individuals with a high nAB value (supplementary figure 11). Genome-wide (coding regions and non-coding regions) we identify 16951 common variants (>5% MAF in at least one population) in the 1000 Genomes data, that are significantly associated with nAB and constitute our set of error candidates. Interestingly, these candidates are not distributed randomly over the genome, but are enriched in coding regions, where around 696 variants (~1%) are predicted to be errors (supplementary table 4-5). In comparison, in non-coding regions only a small fraction (<0.001%) were labeled as candidates, even when more variants are sampled to increase power in the prediction (supplementary table 10). To further test the enrichment in coding regions, we used the software *admixture* (Alexander, et al. 2009), which labels individuals by components of ancestry, on variants in coding regions and on all variants genome-wide. Coding regions showed components that corresponded to the grouping of individuals by nAB and with technical features of the samples (supplementary figure 12), while such an effect was not observed for non-coding regions variants, suggesting that variants in coding regions are enriched for error.

### Presence of error candidates in different datasets

We would expect that our predicted errors are shared less often than real variants with datasets produced at high coverage by different technologies. To test this prediction, we calculate how often our candidate variants are found in the genomes of 69 individuals generated by Complete Genomics and compare this number to the sharing of other frequency matched variants from the 1000 Genomes dataset. In coding regions, 85% of the matched variants are found in the Complete Genomics datasets, while only 15% of our candidates are shared (chi-square test p-value<10^−15^). In non-coding regions 80% of variants match, while only 56% of candidates are shared (p-value<10^−15^). This suggests that around 84% of our predicted variants in coding regions and 33% of variants in non-coding regions are more likely due to error, assuming conservatively that Complete Genomics is devoid of errors that are shared with the 1000 Genomes dataset. These proportions increase for lower FDR thresholds and lower allele frequencies (supplementary figures 14-15). For example, only 42 out of 4681 candidates in non-coding regions that have a frequency <1% are present in Complete Genomics, and 98% of these candidates are estimated to be errors; the fraction of true errors is 56%, when considering all variants lower than 10% frequency.

We also assessed whether these errors are unique to the 1000 Genomes dataset or whether they can be found in other large collections of genomes that may contain similar batch effects. We find that 7843 error candidates, out of which 69 occur in coding regions, are present also in the HRC dataset In the GoNL dataset (The Genome of the Netherlands Consortium, et al. 2014) we find 7380 error candidates, of which 32 are in coding regions. These variants are underrepresented in the Complete Genomics dataset (χ-test < 10^−16^ for both GoNL and HRC), although with a proportion of estimated errors lower than those of the full set of candidates (49% and 17 % in the coding and non-coding regions in GoNL; 69% and 26% in HRC). We also note that while we are able to detect some of the systematic errors from the 1000 Genomes in both datasets, the fraction of predicted errors that are present (56% and 0.49% in GoNL and HRC, respectively) is significantly lower than the fraction of variants that were not labelled as error (67% and 80%; p-values<10^−15^ in both cases). For the HRC dataset, which is based on 1000 Genomes data, this suggests that further filtering was efficient in removing a large proportion systematic errors. Consistenly, in the coding regions only 10% of the candidates are present in both 1000 Genomes and HRC datasets, while 91% of the frequency matched coding variants overlap.

### Characterizing the features of predicted errors

Errors may be caused by a variety of technical issues (Chen, et al. 2017; Dohm, et al. 2008; Schirmer, et al. 2015; Wang, et al. 2017). To learn more about the features of error candidates in the 1000 Genomes dataset, we tested several characteristics in comparison to background variants that were randomly selected from the set of all tested variants. We first tested candidates in coding region, and divided the candidates and control variants into synonymous and nonsynonymous sites. Whereas the control set shows an approximatively equal number of non-synonymous and synonymous variants (~48% non-synonymous), error candidates show a much higher proportion of non-synonymous variants (~72%), consistent with the expected fraction of non-synonymous sites given the codon composition of human genes (~72%) (Nei and Gojobori 1986) (fig. 3). Coding region candidates also show a higher proportion of transversions (z-test p<10^−6^), a base composition with a higher proportion of Gs and Cs (z-test p<10^−12^), a propensity to fall within short C or G homopolymer stretches (z-test p<10^−6^), and an excess of heterozygous genotypes in respect to the Hardy-Weinberg equilibrium (Wilcoxon Rank test on control and candidate p-values: p<10^−12^). The last test indicates that erroneous sites are often heterozygous. Allele imbalance, i.e. the unequal representation of sequences supporting the two alleles in a high-coverage sample, is often used as a hallmark sign for erroneous heterozygotes (Li 2014). To test whether this is also true for our candidates we used 25 high-coverage samples from the SGDP panel (Mallick, et al. 2016) that were independently sequenced with the Illumina platform. Coding region candidates that are present among those samples show a strong imbalance (chi-square test p-value <10^−12^), suggesting that sequencing errors or cross-contamination introduce apparent heterozygous sites (fig. 3, supplementary figure 16). For genome-wide candidates, we find some of the same signals, albeit often weaker. Candidates still exhibit a tendency to reside in homopolymer stretches (z-test p<10^−5^), occur more often in heterozygous state (p<10^−12^) and show allele imbalance (p<10^−12^).

**Figure 3.**
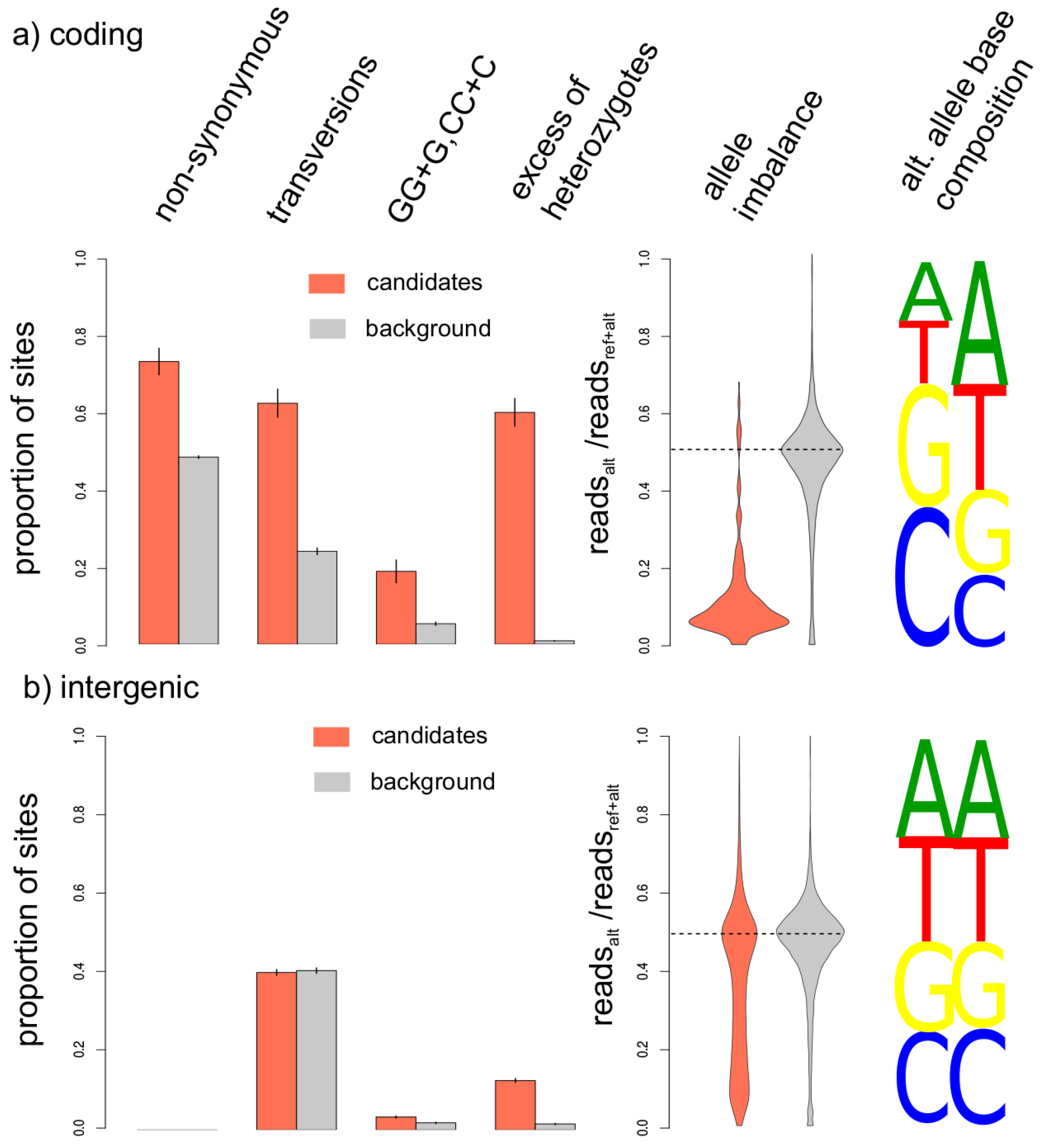
Characteristics of error candidates in the 1000 Genomes dataset detected in coding regions (a) or genome-wide based on the intergenic dataset (b). For error candidates (red) and frequency-matched background variants (gray), the barplot shows the proportions of non-synonymous versus synonymous variants, transversions versus transitions, alternative alleles introducing Gs or Cs after or before GG or CC dimers, and positions with significant excess of heterozygotes (pvalue<0.05). The violin plot shows the proportion of sequences supporting the alternative alleles in individual with at least one sequence showing the alternative allele. On the right, the base composition of alternative alleles is shown for error candidates (red) and background variants (gray).

As part of their release, the 1000 Genomes Project provided users with two annotations, that label variants by high- or low coverage, or the presence of low mapping quality sequences. These two annotations differ in strictness, with one representing more permissive criteria that label fewer bases as potentially problematic (pilot accessibility filter) and the other a stricter filter that labels more variants (strict accessibility filter). We observe that the majority of error candidates in coding regions are not labelled by either annotation, whereas at least 25% remain unlabeled for genome-wide variants (fig. 4). This indicates that, at least in coding regions, inter-chromosomal linkage detects erroneous variants that are not detectable by considering coverage or mappability alone.

**Figure 4.**
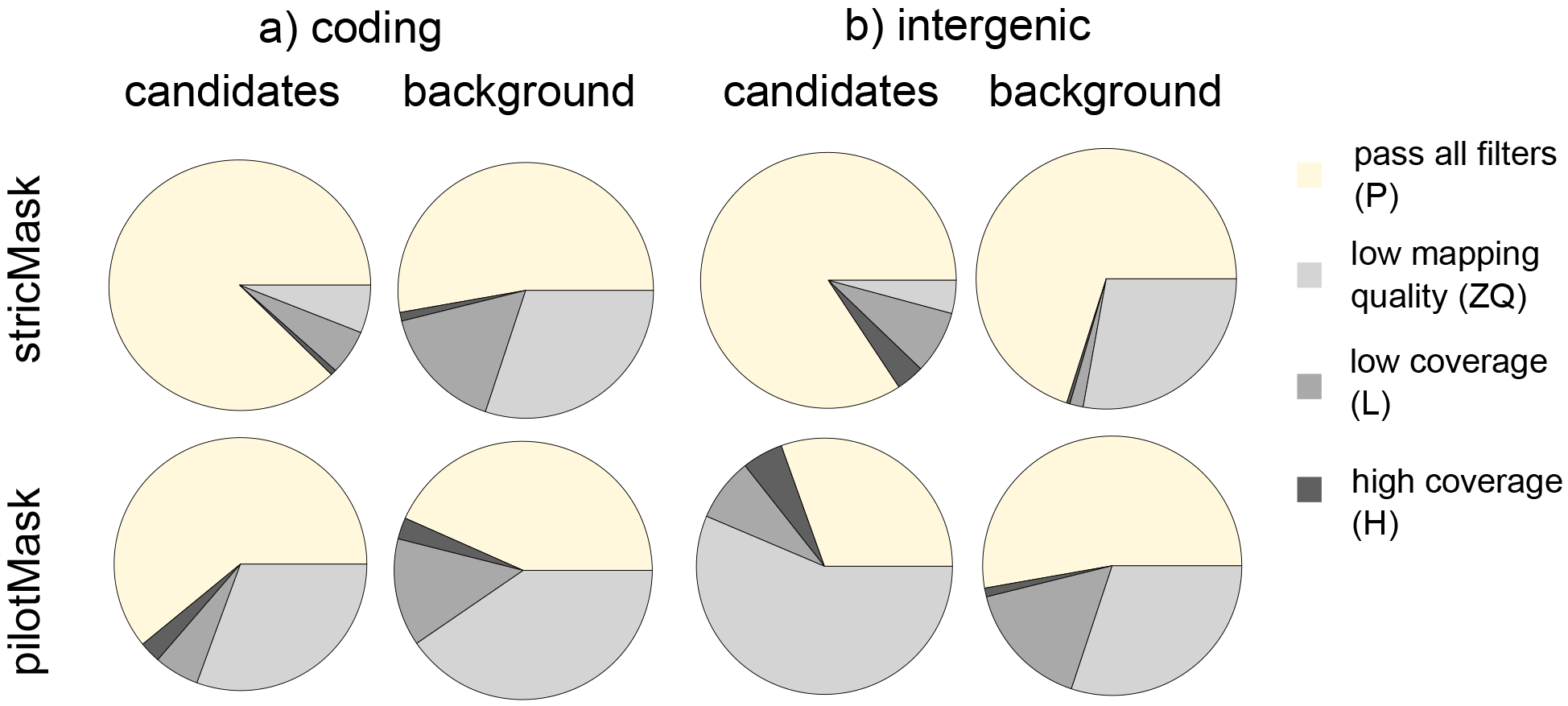
Coverage and mappability features of error candidates (left) and background variants (right) according to two accessibility filters of the 1000 Genomes dataset. The coverage filters exclude regions where the depth of coverage (summed across all samples) was higher or lower than the average depth by a factor of 2-fold (pilot) or by 50% (strict). Regions are deemed as lowly mappable if >20% of overlapping reads have mapping quality of zero (pilot) or >0.1% (strict).

### Testing uniform datasets

Our method predicts errors that likely originate from the combination of technologies, but should not find errors when the dataset is generated with just one technology and does not contain other batch effects. To test this prediction, we analyzed data from the GoNL dataset (The Genome of the Netherlands Consortium, et al. 2014), which is composed of two parts that were produced with the Illumina and Complete Genomics (Drmanac, et al. 2010) sequencing technologies, respectively. We first analyzed the two parts independently and found no excess of linked pairs, consistent with a uniform error within each part. However, when both parts are merged, 2750 linkage pairs with FDR<20% are detected (supplementary figure 17; supplementary table 8), representing batch-specific variants that are likely due to error. We applied our method to detect the variants that drive this signal. Similar to the 1000 genomes dataset the identified variants display typical features of errors, such as an excess of heterozygotes and evidence of allele imbalance (supplementary figure 18).

We also analyzed how many false positive errors we predict in another uniform dataset, containing 654 unrelated individuals from the Botnia region (Fuchsberger, et al. 2016). The dataset was produced as part of a diabetes genome-wide association study using OmniChip. Our method found no excess of linked pairs in this dataset and the distribution of nAB across individuals is comparable to that observed after randomly permuting chromosomes across individuals.

## Discussion

Previous studies used local patterns of linkage disequilibrium (LD) in order to improve the quality of haplotype and SNP-calls in large-scale studies (Leek 2014; Scheet and Stephens 2006). An example is the fastPhase method, that allowed for the identification of over 1,500 low frequency SNPs with high error rates in the HapMap datasets (Scheet and Stephens 2006). Our method uses a different source of information and can be combined with these approaches to predict errors. Here we have shown that long-distance linkage between pairs of sites that reside on different chromosomes can be used to predict individuals that show an excess of error and to label variants that are likely errors.

The errors we detected can influence a variety of analyses. For instance, we showed that they affect estimates of population structure (supplementary figure 12) and estimates of mutational load (supplementary figure 19). Furthermore, since exons are enriched for errors and random errors appear more often as non-synonymous variants, estimates of functional mutational load and the fitness effects of newly arising mutations might be affected. The apparent linkage between chromosomes can also affect studies of epistatic interactions. For example, Sohail et al. were able to detect epistatic effects only for the most functional elements of the genomes, and detected an overall signal of linkage disequilibrium compatible with the presence of errors, as identified in the present study (Sohail, et al. 2017). Our approach allows to identify these errors.

We note that our estimates of the per-individual errors allow for further analyses to study the possible origin of batch effects. For example, we observed in the 1000 Genomes dataset a strong effect of the sequencing center, followed by coverage and genotyping array used (supplementary table 2,3; supplementary figure 8,9). However, the insights from the published meta-information are limited since sequencing center, for instance, could represent a variety of underlying causes for quality differences, such as a differences in chemical reagents or operating conditions (Chen, et al. 2017; Leek, et al. 2010).

Our error candidates showed an excess of heterozygotes. This heterozygotes are characterized by allele imbalance in independently sequenced high-coverage datasets. This suggests that these positions are susceptible to recurrent errors. Note that these errors are elusive, and often not captured by coverage and mappability based filters. We note that our analysis was restricted to variants that passed genotype quality filters (labelled as PASS). However, consistently with the signal of allele imbalance, the genotype quality of these heterozygous errors was on average a bit lower than for other variants (supplementary figure 20).

Inter-chomosomal linkage disequilibrium leverages the usage of different technologies to identify errors that could remain unidentified when only one technology is used for sequencing. We showed this using the GoNL dataset, for which combining two datasets generated with different technologies allows for the discovery of platform-specific errors. Thus, while using a single platform may help in obtaining datasets with errors that are comparable between samples, the combination of these datasets can help identify errors that are different between technologies. We hope that our method helps to increase the value of large-scale heterogeneous datasets that are more susceptible to batch-effects.

## Methods

### Data handling

We downloaded the 1000 genomes phase 3 dataset (version 6/25/2014). One representative individual was sampled for each set of related individuals using the 1000 genomes annotation. Only populations with at least 95 unrelated individuals were analyzed further, retaining 12 populations and 1117 individuals (supplementary table 1). Variants were classified according to frequency using bcftools (common variants: >5% frequency in at least one population, rare variants: 1% < frequency in at least one population, but <= 5% in all) (Li 2011). We performed all analyses on both common and rare variants, or only on common. Variants were annotated as coding when they fell within the coding exons of the UCSC known gene annotation (Rosenbloom, et al. 2015), and as intergenic when they did not overlap UCSC known genes and were not annotated as a potentially functional variant by the Variance Effective Predictor (McLaren, et al. 2016). The Botnia data include 327 trios from the Botnia population, in Finland (Fuchsberger, et al. 2016). We excluded all offspring and related individuals.

Data from the Genome of the Netherlands were filtered and annotated analogously to the 1000 genomes. All analyses shown refer to variants with a 5% MAF cutoff.

### Outline of pipeline

We implemented our analyses in a pipeline to detect inter-chromosomal linkage disequilibrium and detect variants affected by batch effects or inhomogeneity in the treatment of samples. This pipeline is outlined in supplementary figure 1, and the different steps are described in the following sections.

#### Step-1: Linkage Disequilibrium

When the phase is unknown, as for two physically unlinked loci A and B with possible alleles A-a and B-b, respectively, a composite genotypic linkage disequilibrium can be calculated, by relying on a maximum likelihood estimate of the amount of AB-gametes that are present in samples. Following Weir (Weir 1996), we can arrange the counts of the nine possible observed genotypes for the two loci in a matrix:

| A \ B | B/B | B/b | b/b |
| --- | --- | --- | --- |
| A/A | n1 | n2 | n3 |
| A/a | n4 | n5 | n6 |
| a/a | n7 | n8 | n9 |

so that

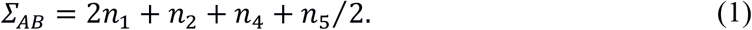

The composite genetic disequilibrium equals *D*_*AB*_ = *Σ*_*AB*_/*n-2p*_*A*_*p*_*B*_ where *n* is the number of samples. The sign of the composite linkage disequilibrium *D*_*AB*_ indicates whether alleles A and B (or a and b) occur preferentially in combination (*D*_*AB*_>0) or whether the alleles occurring most often together are A and b (or a and B) (*D*_*AB*_<0). We can test statistical association between two variants by either considering a two-tailed test (i.e. Fisher’s exact test, adopting normalization proposed by Kulinskaya and Lewin (Kulinskaya and Lewin 2009), or by performing a 1-tailed Fisher’s exact test for the positive and negative associations between minor alleles, thus denoted as A and B.

In order to speed up calculations approximate p-values were first determined with the chi-square based T2 method (Schaid 2004; Wu, et al. 2008; Zaykin, et al. 2008), and exact p-values were calculated only for those pairs with an approximate p-value < 100/nSNP^2^, where nSNP is the total number of variants examined. While negative association between variants might also occur because of synergistic interaction between deleterious variants (Sohail, et al. 2017), batch effects are expected to result in the positive association between rare variants (Fig1). Thus, we restricted our analyses to significantly linked variants with a positive association (but see supplementary figure 21 for an analysis of negative linkage, and supplementary figure 22 for both positive and negative linkage together).

With the exception of the per-population analysis shown in fig. 2a-b, supplementary figures. 6a,b and supplementary figure 10c we combine p-values across populations, using Fisher’s method to obtain a single combined 1-tailed p-value for each pair of variants. These combined p-values are then compared to those obtained in a dataset generated by randomly redistributing chromosomes across individuals. This allows to additionally control for the sporadic linkage between chromosomes that can occur for low frequency alleles (Skelly, et al. 2016). The False Discovery Rate was calculated as the fraction of allele-pairs that have an equal or lower p-value in the randomized dataset, versus the original data. In order to test the excess of inter-chromosomal linkage disequilibrium we restrict further analyses to instances in which at least one pair of variants is significantly associated (supplementary figures. 2-3).

#### Step2: Individual contribution to LD

We calculated the contribution of each sample/diploid individual to the linkage signal by summing up its contribution to the total of the *Σ*_*AB*_ values over all significant linkage pairs. Note that this value, called nAB, is calculated per individual. For positive associations (D>0), the contribution of each sample is directly the weight in equation (1) corresponding to a specific genotype configuration, e.g. 2 if a sample has genotype n1 (A/A, B/B) since both gametes necessarily had alleles A and B on both chromosomes, and 0 if it has genotype n3 (A/A, b/b), since no gametes had alleles A and B on both chromosomes.

In order to test whether samples contribute uniformly to the linkage signal or not we compared the variance of the observed data and random reshuffling of the chromosomes across samples and within populations. To identify samples with similar features, we perform a finite mixture analysis (R package Mixtools), by assuming a normal distribution for the underlying model of the contribution to the linkage signal of each group of samples, within a population. We calculated the corrected Akaike Information Criterion (AICc) weights for each model, with 1 to 9 possible underlying Gaussians for each population.

The models with the highest weights are shown in supplementary figure 7. Since a Gaussian might deviate from the real underlying distribution, we tested whether a finite mixture analysis on the null datasets in which chromosomes are redistributed across individuals would provide less support to the presence of groups of samples with different nAB. We first calculated the relative support for the best supported model against the null model with only one Gaussian to explain the data, and compared it with the same statistics for 100 null datasets. Higher support for multiple clusters was present in the observed data compared to the null distribution (Wilcoxon rank test, p-value<10^−16^ for coding regions, p-value=0.00103 for intergenic). Note that although generating permutations of the data is computationally expensive, the high number of potential links give a very narrow distribution of all statistics related to this empirical null distribution. For the 1000 genomes dataset 3 randomizations are sufficient to provide highly significant p-values when adopting a one-tailed t-test and comparing to the real data. When a dataset showed a significantly higher variance and higher clustering than its empirical null distribution in terms of nAB, we proceed into identifying the variants responsible for the signal and the sources of the bias. In order to assess whether the identified clusters correspond to specific features of the samples we tested the role of several technical predictors extracted from the sample spreadsheet of the 1000 genomes dataset (ftp://ftp.1000genomes.ebi.ac.uk/vol1/ftp/technical/working/20130606_sample_info/20130606_sample_info.xlsx). The clusters identified with Mixtools for the coding regions of the 1000 genomes are significantly associated with Sequencing Center and average coverage (combined chi-square p-value<10^−16^) (supplementary figures 8-9).

#### Step 3: Identification of error candidates

In order to identify variants whose presence is explained by the occurrence in specific clusters of samples, we used a Generalized Linear Mixed Model iteratively on each variant, considering the contribution of each sample to nAB as only predictor. The underlying assumption is that samples that show a consistent excess of linked variants are more likely affected by technical artifacts. Hence, variants that are present only in these samples would be more likely spurious. To assess whether the presence of an allele is predicted by the contribution of each individual to the linkage signal, we built two models: a full model, including the nAB value for each individual as a predictor, and a null model, in which nAB is not included. We then compared the two models with a likelihood ratio test, so that for each variant we assess the significance of the relationship between nAB and the presence of the minor variant (supplementary figure 11). Notice, that this method uses as only predictor the observed linkage, and thus does not require any additional information about the samples. In more detail, the response variable of the linear models is the presence or absence of the minor allele per sample. A sample can have three states for this minor allele (absent homozygous a/a, heterozygous a/A or present homozygous A/A). To model non-independence of the two chromosomes of each individual we consider each allele separately and introduce a predictive variable (factor) “Sample” that groups the two alleles for each chromosome of each individual (supplementary figure 11). The other predictor that is present in both the full and the null model is the population to which each individual belongs. In the full model, the contribution to the linkage signal per sample (nAB), is also present as a continuous covariate.

We can thus write the two models as:

- full model: presence allele A at site i ~ contribution nAB + (1| Population) + (0 + contribution nAB | Population) + (1|Sample).
- null model: presence allele A at site i ~ (1| Population) + (1|Sample).

Sample and Population were introduced as two random factors (categorical random predictors), in order to control for the non-independence between chromosomes belonging to the same individual and individuals belonging to the same population. The effects of random factors are denoted as (1|Factor). The effects of a covariate, when dependent on a random factor, are denoted as (0 + covariate | Factor). In particular, we allowed for different effects of nAB in different populations (0 + contribution nAB | Population), due to potential differences in population composition and treatment.

The contribution of each sample to the nAB signal has been z-transformed and the p-values of the likelihood ratio test are corrected for multiple testing with the Benjamini-Hochberg criterion. In order to speed up calculations we preliminarily scanned each sample with an analogous simpler logistic model, in which random factors are neglected, and populations are considered independently. The p-values of each population are combined with Fisher’s method, and the full model including all random components was performed only for variants for which the combined p-value was below 0.01.

### Analysis of the 1000 Genomes dataset

#### Characterization of batch-effects

To directly assess the association between nAB and technical features of the samples, we applied a Generalized Linear Mixed Model using the R package lme4 (supplementary tables 2-3). We tested a model exploring whether the observed log(nAB) value for each sample (response variable) is predicted by technical features of the individual samples (predictor variables). As predictor variables, we used technical features of the samples described in the sample spreadsheet of the 1000 Genomes project. For simplicity, we grouped these different predictors into three main groups: Center, Coverage and Chip. As a first predictor variable, we considered the main sequencing center where each sample was processed, i.e. for coding regions the main sequencing center for the exome, and for intergenic regions the main sequencing center for the low coverage sequencing. For most samples, one sequencing center was used to produce all or at least the majority of the data, which was regarded as the main sequencing center; for the remaining cases (n<3 for all populations), where equal proportions of data were produced at multiple sequencing centers, we considered this combination as an independent level. Center is a single categorical variable, in which the different levels of the linear model indicate different sequencing centers, and the coefficients estimated by the linear model (supplementary tables 2-3) are the effect that each sequencing center has in respect to a baseline sequencing center selected from the spreadsheet. The second group of variables, Chip, includes three independent binary variables, each denoting whether one the genoyping array platforms (Omni, Affymetrix or Axiom) was used for the sample. Finally, the group Coverage, describes the average coverage per sample, measured as three continuous variables from the sampled spreadsheet of the 1000 genomes dataset, i.e. Total.Exome.Sequence, X‥Targets.Covered.to.20x.or.greater and LC.Non.Duplicated.Aligned.Coverage. Populations were included as random categorical predictors, and for all other predictor variables we considered random intercepts and random slopes nested within Population. This approach accounts for the different effects that the different predictor variables might have in different populations. We tested a full model, that included all predictor variables, and three reduced models including only some of the predictors: 1) the three continuous coverage variables (Coverage) + Center, 2) Coverage + the presence of genotyping arrays (Chip), 3) Center + Chip. The models were compared with a likelihood ratio test, indicating whether the group removed in the reduced model improves significantly the predictions of the model (supplementary tables 2-3).

#### Idenfication of error candidates

We applied our method to all variants present in the 1000 genomes datasets. For the coding regions, where linkage pairs are abundant, we directly use the nAB values estimated exclusively on significantly linked variants using a minimum allele frequency threshold of 5% (supplementary table 4) and 1% (supplementary table 5). This procedure is under-powered for intergenic variants, where the amount of linked pairs is much smaller and the distribution of nAB has a low resolution. To increase the amount of bona-fide linked variants in the intergenic dataset, we first increased by 10-fold the number of pairwise inter-chromosomal comparisons by subsampling a larger amount of intergenic variants. In addition, we relax the FDR cutoff to define linked pairs to FDR<20%. Notice that while this reduced cutoff may increase the noise in the nAB profile due to additional randomly linked pairs, we expect no systematic bias that would increase the number of predicted errors. The set of discovered variants is reported in supplementary table 6 and supplementary table 7, for minor allele frequency cutoff of 5% and 1%, respectively. In contrast, increasing the FDR cutoff for links considered to compute nAB has only a minimal effect on the number of predicted errors in coding regions, suggesting that the estimation of the nAB profile for the exome is not under-powered.

For both datasets, we estimated the false discovery rate for each variant with the Benjamini-Hochberg method (supplementary tables 4-7). An empirical false discovery measure can be obtained by calculating the overlap of the candidate variants from the 1000 genomes dataset to variants present in Complete Genomics (supplementary figs. 14-15). Significant variants for both datasets show a reduced overlap with the high quality Complete Genomics dataset (pvalue<10^−16^), indicating an enrichment in error among our candidates.

### Effects of selection

In order to illustrate the possible selection scenarios that could lead to inter-chromosomal linked variants we calculated the dynamics in time of the average linkage-disequilibrium coefficient D, in presence of epistatic interactions between two different genomic variants leading to a difference in survival rates of the different gametes. We consider two biallelic sites, with alleles A-a, and B-b, respectively. We denote the number and selection coefficient of gametes AB,Ab, Ab and aB with n_AB_,n_Ab_,n_aB_ and n_ab_, and s_AB_,s_Ab_,s_aB_ and s_ab_, respectively. We performed for each selection scenario 10000 simulations. In each generation, we first simulated recombination, then selection. In the recombination step, we sampled the number of gametes that would change state (i.e. gametes AB recombining with ab, and Ab recombining with aB) after random pairing of the gametes and recombination occurring with probability r. The selection step follows a Wright-Fisher model, with each gamete having fitness 1+s_AB_,1+s_Ab_,1+s_aB_ and 1+s_ab_, respectively. We consider two possible scenarios: advantageous combinations of minor alleles, with s_AB_>0 and s_Ab_=s_Ab_= s_ab_ =0, and antagonistic combinations of minor and major alleles (with s_AB_=s_ab_=0 and s_Ab_=s_aB_<0). Selection coefficients were either fixed to 1% (strong selection) or sampled from a distribution of selection coefficients estimated for non-synonymous variants in the human genome (Racimo and Schraiber 2014). Simulations are shown in supplementary figure 5.

### Validation of the methods

We tested the current pipeline on simulated datasets with either 50, 100 or 200 unrelated individual genomes of equal length to that of the coding regions analyzed for the 1000 Genomes dataset. The genotypes of these datasets were randomly sampled from the 1000 Genomes datasets. Each chromosome was sampled independently from the others, to obtain datasets with no residual linkage due to population structure nor errors. The individuals were divided into two batches, one with errors and one without errors. The error-containing batch encompassed either 20% or 50% of individuals. Errors were added to either 10% or 50% of the individuals of the error-containing batch at 0.1% of the sites. Errors were added in the form of false heterozygotes, leading to overall error rates equal between 10^−5^ and 0.000125 per site and individual. Results are shown in supplementary figures 23-24.

## Code and Data Availability

We implemented the genome-wide scan in C, and the logistic models adopted for the identification of biased SNPs in R. The code and tables with the predicted erroneous variants in the 1000 Genomes are available at http://bioinf.eva.mpg.de/LDLD/.

## Acknowledgements

We acknowledge the Genome of the Netherlands Consortium for providing access to the GoNL dataset. We thank Roger Mundry for helpful discussions and two anonymous reviewers for helpful comments.

## Supporting Information Legends

### Supplementary table 1

List of the individuals included for the analyses of the 1000 Genomes dataset.

### S1 Appendix

Supplementary table 2,3 and Supplementary Figures 1-24.

### Supplementary table 4

List of inferred errors for the coding regions with a minimum allele frequency threshold of 5%. The list is in BED format. The optional fields indicate with “pvalue” the p-values of Generalized-Linear-Mixed Models (glmm) or General Linear Model (glm) of either all populations (“all”), or only of those in which the variants is above the frequency threshold. The False Discovery Rate is reported in the fdr fields, named after the corresponding model in the pvalue fields.

### Supplementary table 5

List of inferred errors for the coding regions with a minimum allele frequency threshold of 1%. The list is in BED format. The optional fields indicate with “pvalue” the p-values of Generalized-Linear-Mixed Models (glmm) or General Linear Model (glm) of either all populations (“all”), or only of those in which the variants is above the frequency threshold. The False Discovery Rate is reported in the fdr fields, named after the corresponding model in the pvalue fields.

### Supplementary table 6

Genome-wide list of inferred errors inferred from inter-chromosomal linkage disequilibrium in intergenic regions, with a minimum allele frequency threshold of 5%.

The list is in BED format. The optional fields indicate with “pvalue” the p-values of Generalized-Linear-Mixed Models (glmm) or General Linear Model (glm) of either all populations (“all”), or only of those in which the variants is above the frequency threshold. The False Discovery Rate is reported in the fdr fields, named after the corresponding model in the pvalue fields.

### Supplementary table 7

Genome-wide list of inferred errors inferred from inter-chromosomal linkage disequilibrium in intergenic regions, with a minimum allele frequency threshold of 1%.

The list is in BED format. The optional fields indicate with “pvalue” the p-values of Generalized-Linear-Mixed Models (glmm) or General Linear Model (glm) of either all populations (“all”), or only of those in which the variants is above the frequency threshold. The False Discovery Rate is reported in the fdr fields, named after the corresponding model in the pvalue fields.

### Supplementary table 8

Genome-wide list of inferred errors inferred from inter-chromosomal linkage disequilibrium in the GoNL dataset. The list is in BED format. The optional fields indicate with “pvalue” the p-values of Generalized-Linear-Mixed Models (glmm) or General Linear Model (glm) of either all populations (“all”), or only of those in which the variants is above the frequency threshold. The False Discovery Rate is reported in the fdr fields, named after the corresponding model in the pvalue fields.

